# RNY1 partitions into extracellular vesicles and ribonucleoprotein particles during airway inflammation to regulate macrophage programming

**DOI:** 10.64898/2025.12.04.691704

**Authors:** Cherie E. Saffold, Antiana C. Richardson, Heather H. Pua

## Abstract

YRNAs are small noncoding RNAs that are abundant in both cells and biofluids. Prior research has shown that the secretion of extracellular YRNAs (exYRNAs) changes in response to inflammatory stimuli. However, the mechanisms by which exYRNA packaging and dynamics in biofluids regulate inflammation remain poorly understood. In this study, we found that one YRNA species, RNY1, increased in airway fluid during allergen-induced lung inflammation and correlated with neutrophil infiltration. Using RNase sensitivity assays and size exclusion chromatography, we determined that RNY1 was present in airway fluid extracellular vesicles (EVs) and ribonucleoproteins (RNPs), while another YRNA species, RNY3, was present only in EVs. Both the EV and RNP-containing fractions of airway fluid had a unique ability to program cellular inflammation. Airway fluid EVs increased expression of an alternative activation program in macrophages including *Arg1*, *Ym1*, *Il10*, and *Il6,* while RNPs induced gene expression more consistent with a classic pro-inflammatory phenotype. RNY1 contributed to the programming of macrophages by airway EVs, as macrophages treated with EVs isolated from *RNY1^-/-^* mice demonstrated lower induction of *Arg1* and *Ym1.* Together, these results define the form and function of exYRNAs in lung biofluid and support their role in communicating signals during inflammation.

## Introduction

Extracellular RNAs (exRNAs) are present in all biofluids and have garnered widespread interest due to their potential as diagnostic biomarkers.^1–4^ Growing evidence also points to critical roles for exRNAs in cell-to-cell communication;^5–9^ however, the packaging, transport, and function of individual exRNAs *in vivo* remains poorly understood. While many classes of RNAs are secreted into the extracellular space, exRNAs are enriched for small noncoding RNAs that are <200 nucleotides in length and play critical regulatory roles in the cell. The best studied RNAs in this class are microRNAs (miRNAs), which are selectively secreted and capable of regulating gene expression post-transcriptionally in recipient cells.^10–21^ Yet, miRNAs comprise only a small fraction of all exRNAs.^4,10,22^ The secretion and function of other exRNA species remains a critical gap in our understanding of intercellular communication via RNA.

Among the most commonly detected exRNAs are YRNAs, which are secreted by various cells including cancer cells, immune cells, and stem cells.^10,23,24^ They are highly conserved, with YRNAs or YRNA-orthologs identified in a wide variety of species from vertebrates to bacteria and archaea.^25,26^ The two murine YRNAs (RNY1 and RNY3) are broadly expressed RNA Polymerase III transcripts that exist as ∼110 nucleotide stem-loop structures. Although our understanding of YRNA function remains limited, our best evidence suggests that within the cell, YRNAs bind to protein complexes that control the degradation of other intracellular RNAs.^27^ In addition, YRNAs can be cleaved into smaller fragments,^28,29^ though the regulation and function of these cleavage events are not well understood.

Like other exRNAs, YRNAs are commonly identified in biofluids including plasma, urine, and saliva.^4,30,31^ Within the extracellular space, extracellular YRNA (exYRNA) levels are dysregulated in the blood of patients with heart disease, cancer, and endotoxemia.^32–34^ Furthermore, cells exposed to immune-activating stimuli such as LPS can significantly alter their secretion of YRNAs.^35,36^ But exYRNAs appear to be more than just passive biomarkers of disease, as they have been shown to have immunoregulatory properties. YRNAs from extracellular vesicles (EVs) and immune complexes can induce cytokine secretion, NF-κB pathway modulation, and cell death in recipient cells.^36–40^ These findings support that exYRNAs can instruct inflammatory cell programming, but how exYRNA packaging in biofluids contributes to tissue pathology during disease is incompletely understood.

In this study, we used murine models of airway inflammation to investigate the role of exYRNAs in asthma-like immune responses. We tested (1) how YRNA levels change locally in the lung during tissue inflammation, (2) how exYRNAs are packaged in airway fluid, (3) how the packaging of exYRNAs change with airway inflammation, and (4) how YRNAs contribute to the immunomodulatory properties of vesicles and complexes in the extracellular space. By defining the packaging and immunological roles of extracellular YRNAs, this study reveals both heterogenous packaging of YRNA species and a mechanism by which exYRNAs can shape innate immune responses at local sites of tissue inflammation.

## Methods

### Animals

All animal studies were performed after obtaining approval from the Vanderbilt Institutional Animal Care and Use Committee. RNY1 knockout mice were obtained from GemPharmatech (strain T049875). All other mice were wildtype C57BL/6J (JAX 000664). Mice were used between 7-16 weeks and were housed in a specific pathogen free facility. All airway challenges and intraperitoneal injections were performed under isoflurane anesthesia.

### Ovalbumin (OVA) airway inflammation model

A stock solution of 10 mg/mL OVA protein from chicken egg white (Millipore Sigma A5503) was prepared by resuspending 50 mg OVA in 5 mL pharmaceutical-grade PBS (VWR K812). For sensitization via intraperitoneal injection, 50 μg of OVA per mouse was used. 5 µl of 10 mg/mL OVA was combined with 95 µl pharmaceutical-grade PBS and 100 µl Imject Alum (ThermoFisher 77161) for 200 μL total volume per mouse. This solution was incubated for 0.5-1 hour at room temperature with rocking. Mice were intraperitoneally injected with 200 μL OVA+alum solution on day -9. For airway challenge, 50 μg of OVA per mouse was used. 5 µl of 10 mg/mL OVA was combined with 15 μL pharmaceutical-grade PBS for 20 μL total per mouse. Mice were challenged by oropharyngeal aspiration of 20 μL OVA for allergic airway mice or 20 μL PBS vehicle control on days -2, -1, and 0. Mice were euthanized 1, 4, 9, or 13 days after the last challenge.

### House dust mite (HDM) airway inflammation model

A 100 mg/mL stock solution of HDM extract (Greer XPB70D3A25) was prepared by resuspending 159.7 mg extract in 1.597 mL pharmaceutical grade PBS (VWR K812). A 10 mg/mL working stock of HDM extract was prepared by resuspending 40 μL stock solution in 360 μL pharmaceutical grade PBS. For oropharyngeal aspiration, 40 μg HDM extract per mouse was used. 4 μL of 10 mg/mL HDM extract was diluted in 36 μL pharmaceutical grade PBS, for 40 μL total volume per mouse. Mice were challenged by oropharyngeal aspiration of 40 μL HDM extract for allergic airway mice or 20 μL PBS vehicle control. Mice were challenged on days -16, -14, -12, -9, -7, -5, -2, -1, and 0. Mice were euthanized one day after the last challenge.

### Airway cell and exRNA collection

Euthanized mice were cut in the upper abdomen to expose the diaphragm. An 18g needle was used to pierce the diaphragm as a secondary method of euthanasia. Mice were then cut peripherally towards the head to expose the trachea. A small incision was made in the trachea to insert a catheter. For RNA collection, the lung was slowly washed with 1 mL of ice-cold PBS to collect bronchoalveolar lavage fluid (BALF). For BALF cell flow cytometry, the lung was slowly washed with 1 mL of ice-cold FACS buffer (PBS + 2% FBS + 0.02% sodium azide) to collect BALF. Washing was repeated 2-4 times. BALF washes were then spun at 500xg for 5 minutes to pellet BALF cells. The BALF cell pellet was used for downstream processing. The supernatant was decanted and spun at 2000xg for 30 minutes to remove cell debris. The supernatant was decanted again and used for downstream processing.

### Lung tissue RNA collection

After BALF collection, lungs were excised from the mouse and placed in FBS-containing media. Lungs were then washed with PBS. A ∼50mg piece of lung was cut off, minced, and placed in a tube containing 1 mL of Trizol (Invitrogen 15596026). 500 μL of 1mm soda lime glass beads (Biospec products) were added to each tube. Tubes were beaten in a Mini Beadbeater (Biospec products) for 1 minute to homogenize lung tissue. The resulting Trizol/lung tissue mixture was placed in a fresh tube for RNA extraction.

### RNA extraction

All solid samples (cells/tissues) were placed in Trizol Reagent (Invitrogen 15596026). All aqueous samples (BALF, UF100, UF10, SEC fractions) were placed in Trizol LS Reagent (Invitrogen 10296010). RNA was isolated via phenol-chloroform extraction and isopropanol-based precipitation. Glycoblue Coprecipitant (Invitrogen AM9516) was added to each sample to improve RNA pellet visibility.

### Reverse transcription

For noncoding RNAs, RNA was reverse transcribed using the Mir-X miRNA First Strand Synthesis Kit (Takara 638315) according to manufacturer’s instructions. For mRNAs, RNA was reversed transcribed using Superscript IV First Strand Synthesis System (Invitrogen 18091050) according to the manufacturer’s instructions.

### qPCR

qPCR reactions consisted of cDNA, forward/reverse primers (see Supplemental Table 1), and FastStart Universal SYBR Green Master ROX (Roche 04913914001). qPCR reactions were run using the following protocol:1: 95.0°C for 10:00, 2: 95.0°C for 0:15, 3: 61.0°C for 0:30, 3:Plate Read, 4: GOTO 2, 40 more times.

### Flow Cytometry

BALF cells were incubated in ACK lysis buffer to remove red blood cells and counted using a hemocytometer. Cells were then incubated with Fc block (Invitrogen 14-0161-85) at a 1:1000 dilution and Fixable Viability Dye eFluor 780 (Invitrogen 65-0865-14) at a 1:500 dilution. After washing once with FACS buffer, BALF cells were incubated with the following antibodies at a 1:8 dilution: PE-CD11b (Tonbo 50-0112-U100), PerCP Cy5.5-Siglec F (BD Biosciences 565526), PE Cy7-F4/80 (Tonbo 60-4801-U100), FITC-Ly6G (Biolegend 127606), VioletFluor450-CD45 (Tonbo 75-0451-U100), and APC-CD11c (Invitrogen 17-0114-82). Lymph node cells were used for unstained and single stained compensation controls. Flow cytometry was performed using the FACSCanto II flow cytometer (BD biosciences). Data was analyzed using FlowJo software.

### BALF Ultrafiltration

Ultrafiltrates were purified from BALF from PBS-challenged or OVA-challenged mice. BALF was concentrated using an Amicon ultracentrifugal filter with a 100 kDa pore size (Millipore UFC 910024). The retentate (UF100) was used for downstream processing and experiments. The sub-100 kDa flowthrough was concentrated using an Amicon ultracentrifugal filter with a 10 kDa pore size (Millipore UFC 901024). The retentate (UF10) was used for downstream processing and experiments.

### RNase sensitivity assays

250 μL of unconcentrated BALF, concentrated UF100 EVs, or UF10 samples were incubated with 40 ug/mL RNase A (Thermo Scientific EN0531) for 15 minutes at 37°C. Proteinase K treated samples were incubated with 500 μg/mL proteinase K (Fisher Scientific BP-1700-100) for 1 hour at 37°C treatment prior to RNAse treatment. Triton X-100 treated samples were vortexed for 30 seconds in 0.1% Triton X-100 (Sigma T8787) solution prior to RNase treatment. Samples subjected to all three treatments were incubated with Triton X-100, then Proteinase K, then RNase.

### Size exclusion chromatography (SEC)

500 μL of UF100 concentrates were resuspended with PBS to 500 μL and fractionated using a 35 nm qEV Original v2 size exclusion column (Izon) according to the manufacturer’s instructions. After the void volume was discarded, 8 fractions were collected in 400 μL increments. For RNase sensitivity assays and western blots, SEC fractions 1-4 were pooled.

### Nanoparticle Tracking Analysis

The concentration and size of EVs was analyzed by nanoparticle tracking analysis (NTA) using a ZetaView (ParticleMetrix) with the following parameters: Laser wavelength = 488 nm, filter wavelength = scatter, sensitivity = 70, Shutter = 100. EVs were diluted 1000 to 10,000 times in PBS and measurements were performed using 1 cycle and 11 camera positions. Data were analyzed using ParticleMetrix’s NTA software.

### Western blot

Western blots were performed on BALF EVs or the UF10 from OVA-challenged mice. Western blots were performed with the Jess automated western blot system (BioTechne) according to the manufacturer’s instructions for western blot + total protein quantification replex assay. 600 ng of protein lysate was used per sample. Samples were stained with the following antibodies at a 1:20 dilution: anti-TSG101(Abcam AB30871), anti-CD9 (Abcam AB307085), and anti-Calnexin (Abcam AB22595). Data were analyzed using Compass for SW western blot analysis software (Biotechne). The measured chemiluminescent peak area was used for protein detection, quantification, and statistical analysis. All peak areas were normalized to the total protein loaded per lane, which was calculated by Jess software.

### Bone marrow derived macrophage culture

All mice used for bone marrow derived macrophage (BMDM) culture were derived from female mice. Bone marrow cells were isolated from C57BL/6J murine femurs and tibias. Bone marrow cells were cultured for 5 days in 80% FBS-containing RPMI media + 20% L929 fibroblast conditioned media. On day 6, BMDMs were seeded in 96-well plates at 200,000 cells/well and switched to 100% FBS-containing RPMI media. BMDMs were used for experiments on day 7 of culture.

### Macrophage treatments with airway EVs and Ribonucleoprotein particles (RNP)

EV (UF100) and RNP (UF10) enriched BALF fractions for treatments were obtained through ultrafiltration as previously described. EV and RNP fractions were concentrated to 160 μL and stored at 4°C for 5 days. Before treatment, BMDMs were washed 3x with PBS and resuspended in Opti-mem media without FBS. EV-enriched fractions from *RNY1^+/+^* and *RNY1^-/-^* BALF were added to BMDM cultures at equal volume (10 μL EV fraction in 90 μL media) or at equal particle number (4.2×10^9^-5×10^9^ particles/culture). RNP-enriched fractions from *RNY1^+/+^* and *RNY1^-/-^*BALF were added to BMDM cultures at equal volume (10 μL EV fraction in 90 μL media) or at equal protein amount (15-33 ug/culture). BMDMs were collected after 18 hours and put into Trizol for RNA extraction.

### Classical and alternative macrophage polarization

Before treatment, BMDMs were washed 3x with PBS and resuspended in Opti-mem media without FBS. To classically activate macrophages, the following was added to BMDM cultures: 0.5 ng/mL LPS (Thermofisher 00-4796-93), 0.5 ng/mL IFN-γ (Peprotech 315-05). To alternatively activate macrophages, the following was added to BMDM cultures: 50 ng/mL IL-4 (Peprotech 404-ML/CF), 50 ng/mL IL-13 (Peprotech413-ML/CF). BMDMs were collected after 18 hours and put into Trizol for RNA extraction.

## Results

### Extracellular RNY1 levels increase during airway inflammation

Circulating exYRNA levels increase in autoimmune disorders, cancer, and cardiovascular disease.^1–5^ Yet, it is not well understood how extracellular YRNA dynamics change at local tissue sites during inflammation. The lung offers an excellent system to investigate tissue immune responses with an accessible biofluid, which we and others have shown contain exRNAs that fluctuate in response to inflammation.^41–44^ To investigate how exYRNAs change during airway inflammation, we used a model of asthma where mice are first sensitized to the antigen OVA then challenged in the airway to induce a local allergic response **(Fig S1A)**. To quantify levels of YRNAs, we performed qPCR for the two mouse YRNAs (RNY1 and RNY3) in total BALF cleared of cells and debris by serial centrifugation (500xg x 5min, 2,000g x 30min). Primers were designed to capture both full length and fragmented YRNAs, as these small noncoding RNAs have been shown to exist in both forms in the extracellular space **(Fig 1A)**.^34,35,38,45,46^ We observed an ∼8 fold increase in RNY1 levels in mice challenged with OVA when compared to mice challenged with PBS vehicle control. RNY3 and other Pol III-transcribed exRNAs were not significantly increased **(Fig 1B)**. This preferential increase in BALF exRNY1 levels also occurred in HDM-challenged mice, a second model of sterile, allergen-induced lung inflammation **(Fig S1B, Fig 1C)**. To determine if increased RNY1 levels could result from increased cellular RNY1 levels, we quantified this YRNA in BALF cells and lung tissue. Interestingly, there was a significant decrease in total RNY1 expression in BALF cells and a trend toward decreased RNY1 levels in lung tissue with airway inflammation **(Fig 1D,E).** These results show that RNY1 levels in the extracellular space selectively increase during airway inflammation with a concomitant decrease in intracellular YRNA expression.

**Figure 1:**
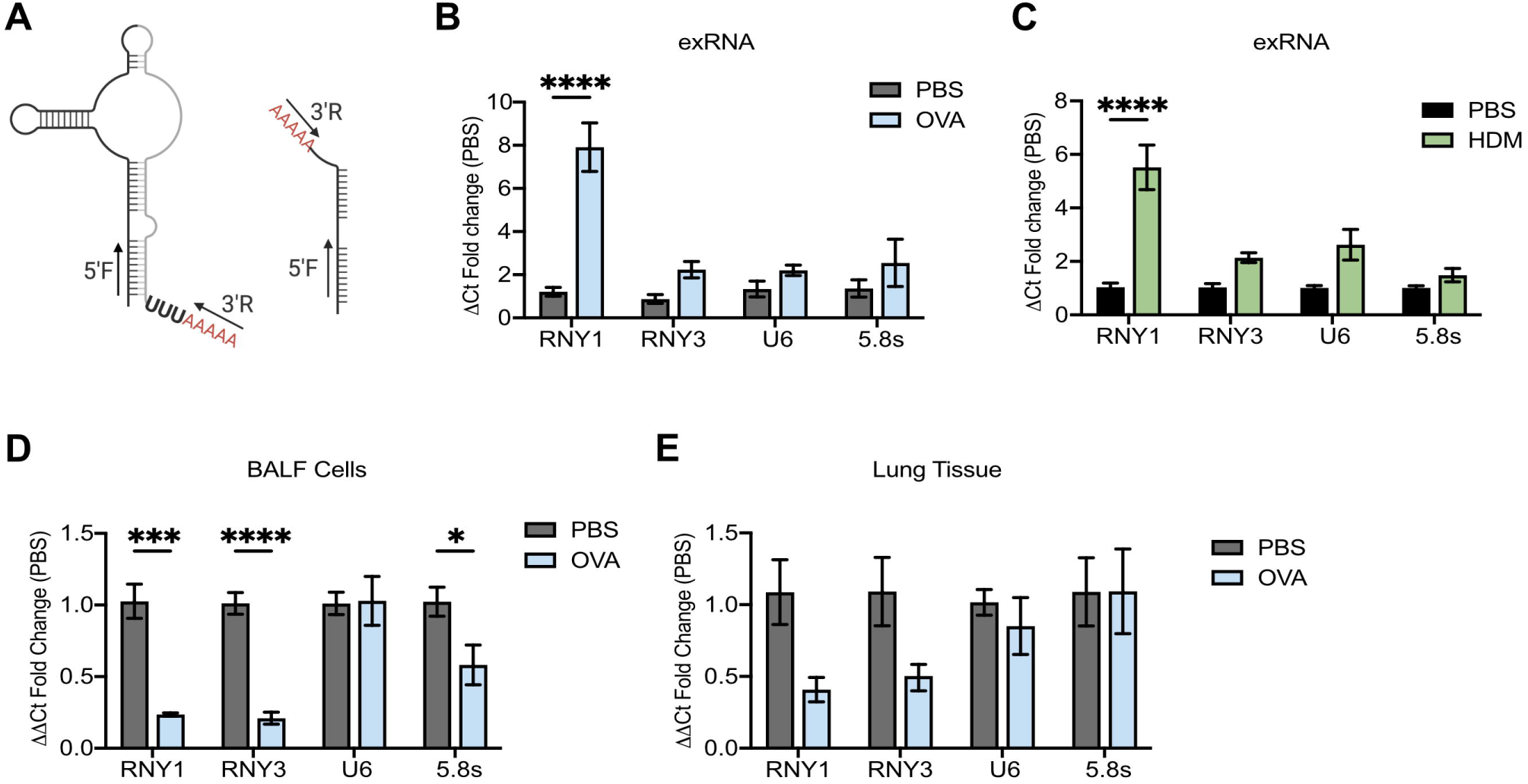
Extracellular YRNA levels increase during airway inflammation. **A)** YRNA qPCR schema. After BALF RNA extraction, RNAs were polyadenylated and reverse transcribed. Each primer set has a primer that detects that 5’ end of the YRNA (5’F) and a universal reverse primer (3’R) that binds to the added polyadenylated tail. The primer sets detect both full length and 5’ fragments of YRNAs. **B-C)** RNA qPCR of murine BALF exRNA one day after the last airway challenge in the OVA **(B)** and HDM **(C)** models of airway inflammation. N= 4-10 from 2+ independent experiments **D-E)** RNA qPCR of murine BALF cells **(D)** and lung tissue **(E)** 1 day after inducing the OVA model of airway inflammation. N= 5 from 2 independent experiments. For all panels: *= P ≤ 0.05, **= P ≤ 0.01, ***= P ≤ 0.001, ****= P ≤ 0.0001. Bars represent the mean +/-standard error of the mean. Two-way ANOVA with Bonferroni.

### Extracellular RNY1 kinetics correlate with immune cell infiltration during airway inflammation

Since we observed an increase in exRNY1 levels during airway inflammation, we wanted to determine how exRNY1 fluctuates with transient and resident immune cell populations in the lung over time. Therefore, we measured exRNY1 and immune cell levels 1, 4, 9, and 13 days after the induction of airway inflammation **(Fig 2A).** RNY1 levels peaked one day after the last airway allergen challenge, then decreased over time **(Fig 2B).** To test whether the levels of RNY1 correlated with specific immune cell populations over the course of inflammation, we measured eosinophil, neutrophil, and macrophage cell dynamics in BALF using flow cytometry **(Fig S2).** We observed a strong significant correlation of extracellular RNY1 levels with neutrophils (R^2^= 0.85), which peaked and then rapidly declined after the last airway challenge **(Fig 2C,D)**. We did not see a significant correlation of RNY1 levels with eosinophils, which peaked 4 days after the last airway challenge **(Fig 2E,F)**. There was no correlation between extracellular RNY1 levels and macrophages, which are a resident immune cell population whose numbers contracted slightly in this inflammatory model **(Fig 2G, H)**. Together, these data show that during inflammation, increases in exRNY1 levels positively correlate with rapidly responding neutrophil infiltration within the local tissue microenvironment.

**Figure 2:**
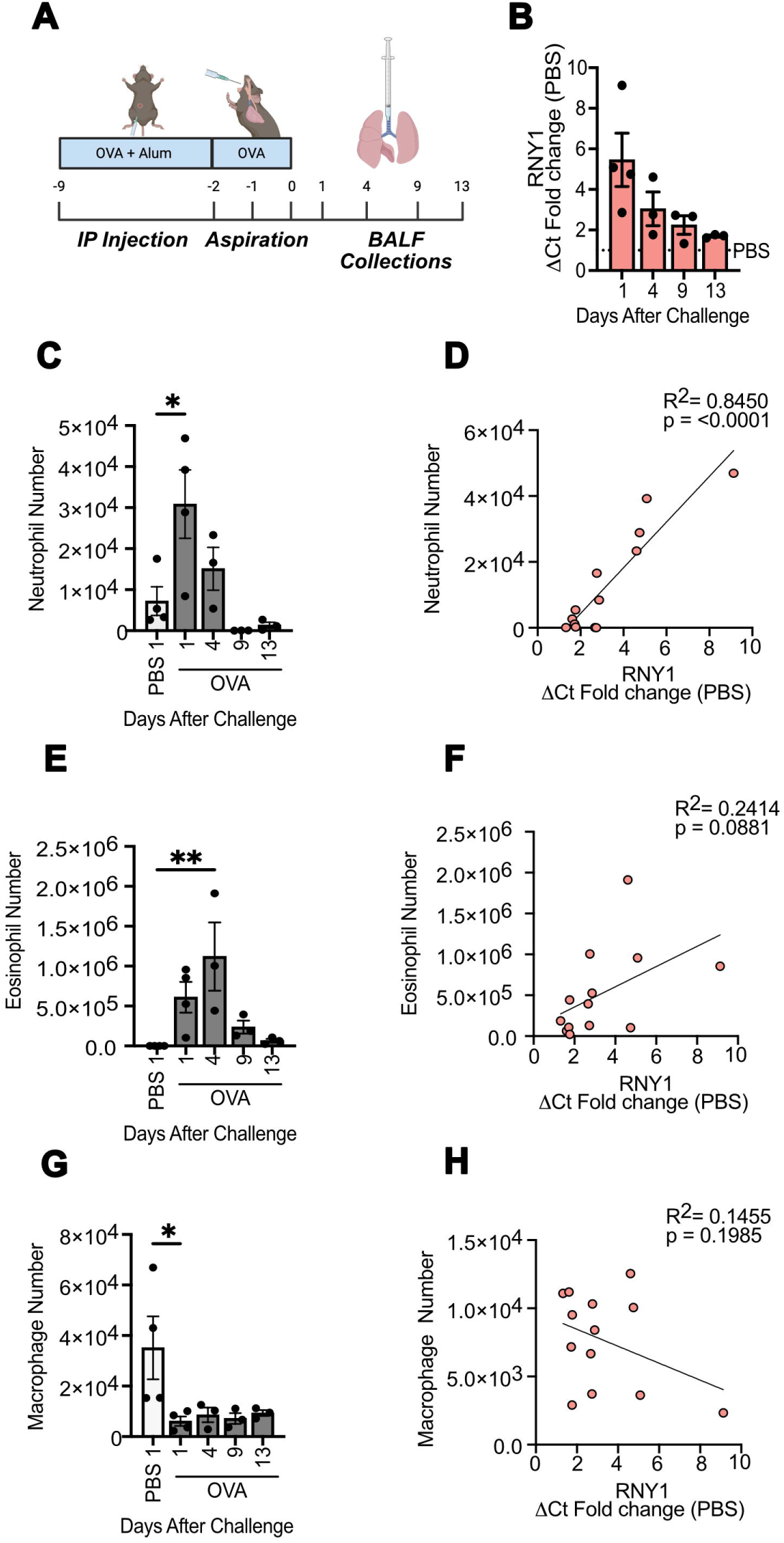
exRNY1 levels correlate with neutrophilic inflammation in the lung. **A)** BALF collection timepoints after OVA airway challenge. **B)** YRNA levels in murine BALF collected 1,4,9, and 13 days after OVA induced airway inflammation. **C,E,G)** BALF neutrophils **(C)**, eosinophils **(E)**, and alveolar macrophage **(G)** counts from murine BALF collected 1,4,9, and 13 days after OVA induced airway inflammation determined via flow cytometry. One way ANOVA with Bonferroni. **D,F,H)** Linear regression analyses of BALF YRNA fold changes vs eosinophil **(D)**, neutrophil **(F)**, and alveolar macrophage **(H)** numbers over time. For all panels: *= P ≤ 0.05, **= P ≤ 0.01, ***= P ≤ 0.001, ****= P ≤ 0.0001. Bars represent the mean +/-standard error of the mean. N= 3-4 mice from each timepoint.

### Extracellular YRNAs increase in extracellular vesicle and ribonucleoprotein compartments during airway inflammation

How RNAs are packaged in the extracellular space is fundamental to understanding their sources, secretion pathways, stability, and function in cellular communication.^47^ To determine whether exYRNAs are stably packaged in the airway, we performed an RNAse sensitivity assay. As expected, pre-extracted BALF was RNAse-sensitive, indicating that naked YRNA sequences are susceptible to RNase degradation. In contrast, ∼75% of exRNY1 and ∼95% of exRNY3 were RNAse-resistant in native BALF at steady state **(Fig 3A)**. Together, these results demonstrate that exYRNAs are protected from degradation by extracellular structures and not by modifications of the RNA itself. Because extracellular RNAs are often packaged inside lipid-bilayer enclosed EVs, we treated BALF with the non-denaturing detergent triton X-100 and evaluated YRNA levels after RNase treatment. Both RNY1 and RNY3 were significantly degraded after detergent treatment, consistent with them being present in lipid-containing EVs. However, ∼25% of RNY1 remained RNase-resistant after detergent treatment **(Fig 3B)**. Therefore, we evaluated whether a subset of RNY1 in the extracellular space was instead protected from degradation by proteins. When BALF was treated with proteinase K + RNase, ∼35% of RNY1 degraded **(Fig 3C)**. All RNY1 was degraded when BALF was co-treated with detergent, proteinase K and RNAse **(Fig 3D)**. Additionally, neither proteinase K nor detergent disrupted the ability of RNAse to degrade YRNAs from extracted BALF RNA, demonstrating that the observed patterns were not the result of interference in enzymatic activity **(Fig S3)**. Finally, we observed no statistically significant differences in RNAse sensitivity patterns in BALF from PBS or OVA treated mice **(Fig 3E-G)**, demonstrating that YRNAs maintain similar compartmentalization patterns at steady state and during airway inflammation.

**Figure 3:**
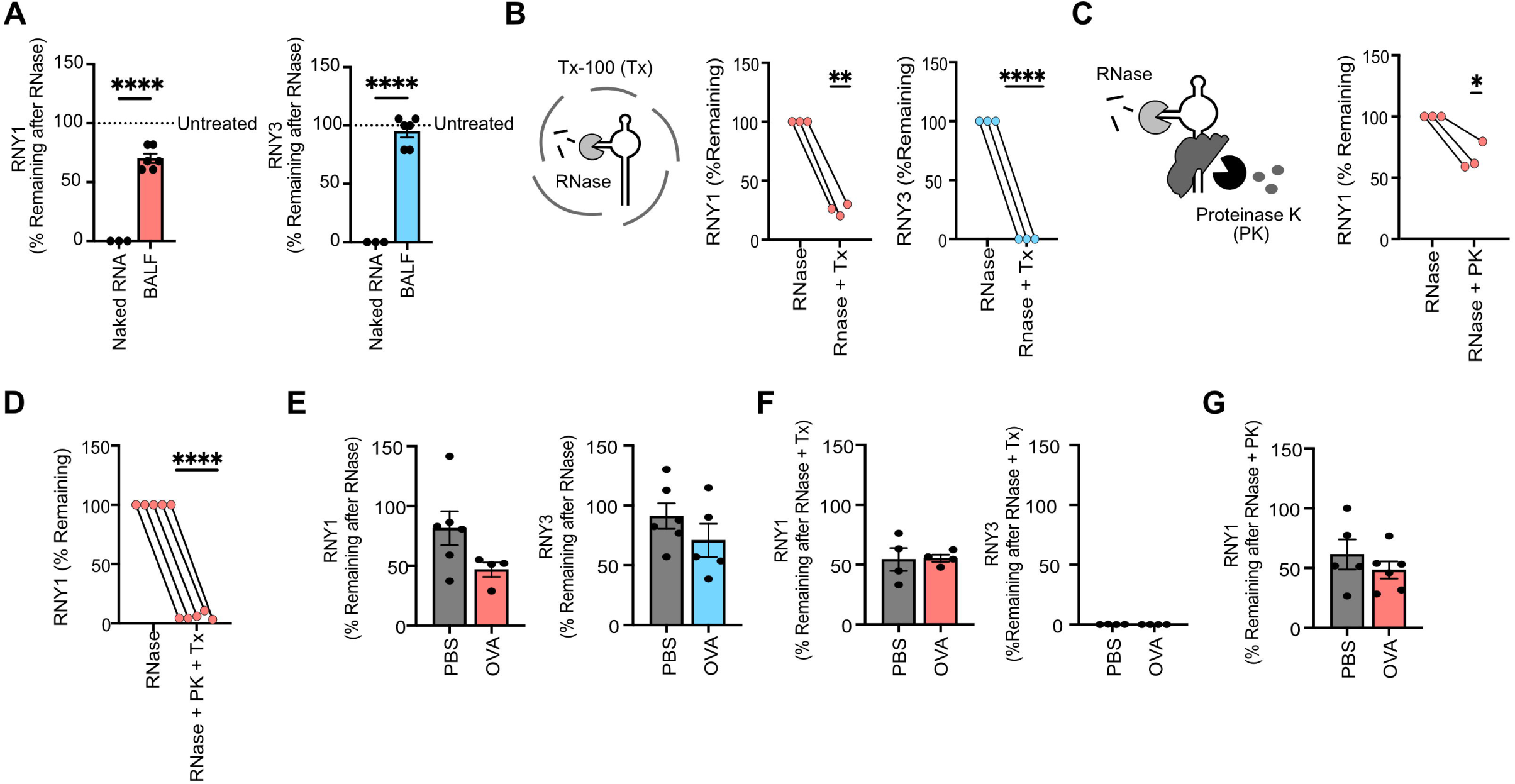
Extracellular YRNA are protected by extracellular vesicles and in protein complexes during airway inflammation. **A)** YRNA qPCR of RNAse-treated BALF. Data is expressed as the percentage of remaining YRNA compared to an untreated control. Naked BALF RNA was pre-extracted via phenol-chloroform. Welch’s t-test. N=3-6 mice from 2-3 independent experiments. One RNY3 value was removed based on a Grubb’s outlier test. **B-C)** YRNA qPCR of Triton X-100 (Tx) + RNase-treated BALF **(B)** and Proteinase K (PK) + RNAse-treated BALF **(C)**. Data is expressed as the percentage of remaining YRNA compared to an RNAse-only treated control. One sample students t-test. N=3 mice from 2 independent experiments **D)** RNY1 qPCR of Tx + PK + RNase-treated BALF. Data is expressed as the percentage of remaining YRNA compared to an RNase-only treated control. One sample student’s t-test. N=5 mice from 2 independent experiments. **E)**YRNA qPCR of RNAse-treated BALF from PBS and OVA-challenged mice. Data is expressed as the percentage of remaining YRNA compared to an untreated control. Unpaired student’s t-test. N= 4-6 from 2-3 independent experiments. One RNY1 value was removed as an outlier using the ROUT outlier test. **F,G)** YRNA qPCR of PK + RNase-treated **(F)** and Tx + RNase-treated **(G)** BALF from PBS and OVA-challenged mice. Data is expressed as the percentage of remaining YRNA compared to an RNase-treated control. Unpaired student’s t-test. N= 4-6 from 2 independent experiments. For all panels: *= P ≤ 0.05, **= P ≤ 0.01, ***= P ≤ 0.001, ****= P ≤ 0.0001. Bars represent the mean +/-standard error of the mean.

To further investigate the heterogenous compartmentalization of exYRNAs in airway fluid during inflammation, we attempted to separate EV-protected and protein-protected YRNAs using differential ultrafiltration. BALF was separated into large (>100Kda, UF100) and small (<100KDA and >10Kda, UF10) fractions **(Fig 4A)**. RNY1 and RNY3 were detected by qPCR in both the UF100 and UF10 fractions. However, we detected ∼16-fold (4 cycles) more RNY1 in the UF100 than UF10 fraction and ∼8,000-fold (13 cycles) more RNY3 in the UF100 than UF10 fraction **(Fig 4B)**. These data indicate that YRNAs are enriched in the UF100 fraction. To test if YRNAs are contained within EVs in the UF100, we performed SEC on this fraction. Both RNY1 and RNY3 cofractionate with EVs, as peak YRNA levels and particle counts were observed in fractions 1-4 **(Fig 4C,D).** Within fractions 1-4, YRNAs were protected from RNase unless treated with detergent **(Fig 4E-F).** As expected, SEC fractions 1-4 were enriched in EV markers including Tsg101 and CD9 **(Fig S4A-C)**. In contrast, eight times more total protein was present in the UF10 fraction than the UF100 fraction **(Fig 4G)**, suggesting that the UF10 is a protein-enriched fraction. In the UF10, ∼65% of RNY1 was RNase resistant, while <1% of RNY3 was RNase resistant, demonstrating that only RNY1 is protected in the UF10 **(Fig 4H)**. All RNY1 in the UF10 was proteinase K sensitive, consistent with RNY1 in this fraction being protected in RNPs **(Fig 4I)**. The same YRNA co-fractionation and RNAse sensitivity patterns were observed in PBS-challenged mice **(S4D-G).** These data show that while protected RNY3 in airway fluid is solely in EVs, RNY1 partitions in both EVs and RNPs with and without airway inflammation.

**Figure 4:**
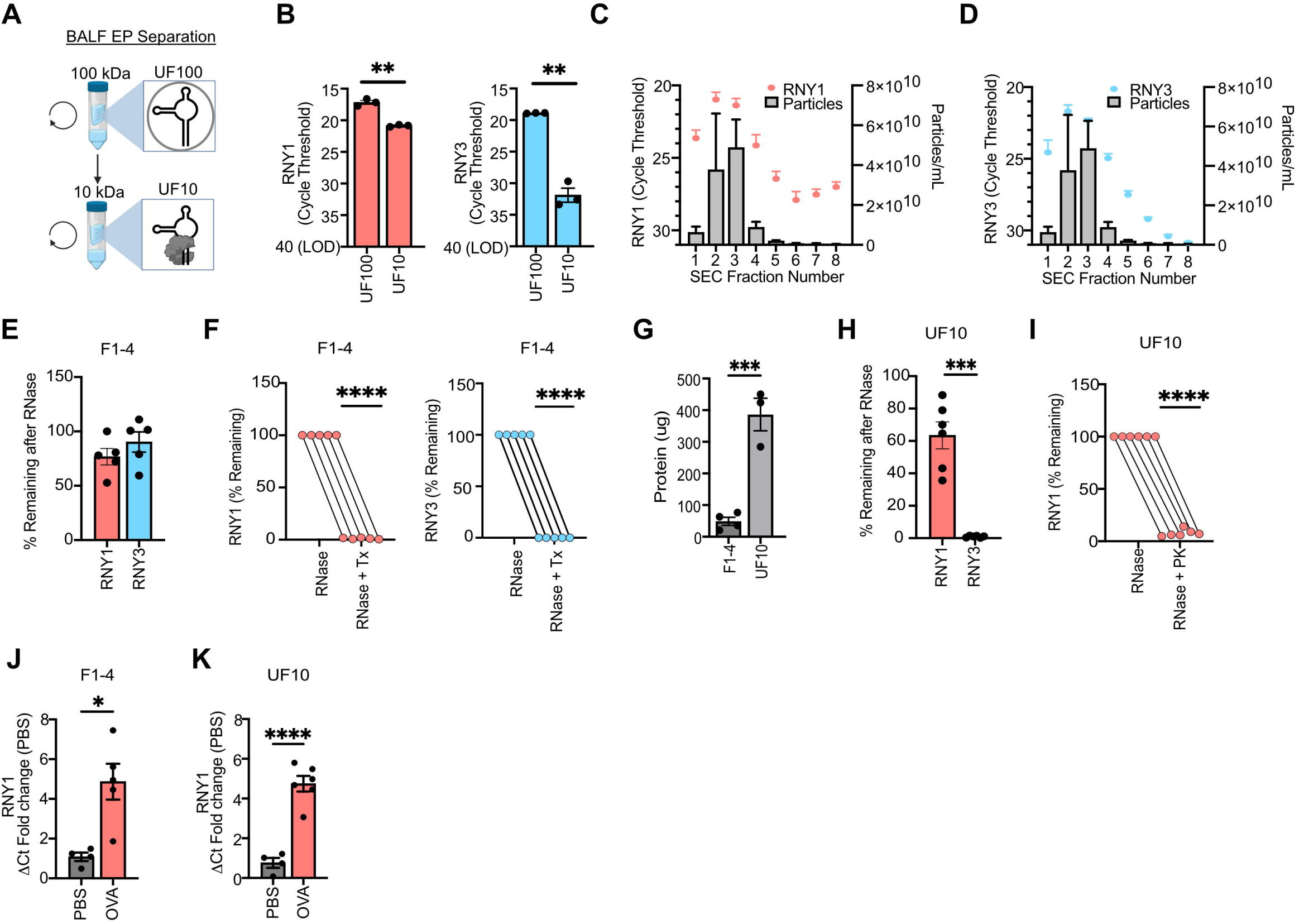
YRNAs partitioned into both extracellular vesicle and ribonucleoproteins increase during airway inflammation. **A)** Schematic of BALF partitioning into UF100 and UF10 fractions. The UF100 fraction contains material > 100 kDa. The UF10 fraction contains material < 100 kDa and > 10 kDa. **B)** Ct values for RNY1 (left) and RNY3 (right) in UF100 and UF 10 fractions. N=3 from 1 independent experiment. Unpaired student’s t-test. **C,D)** RNY1 **(C)** and RNY3 **(D)** qPCR Ct values and particle counts from SEC-fractionated BALF UF100. N=3 from 2 independent experiments. **E)** YRNA qPCR of RNAse-treated BALF SEC fractions 1-4. Data is expressed as the percentage of remaining YRNA compared to an untreated control. N=5 from 2 independent experiments. Unpaired student’s t-test. **F)** YRNA qPCR of Triton X-100 (Tx) + RNase-treated BALF SEC fractions 1-4. Data is expressed as the percentage of remaining YRNA compared to an RNase-only treated control. One sample student’s t-test. N = 5 from 2 independent experiments. **G)** Qubit protein quantifications of BALF SEC fractions 1-4 and UF10. N=3-4 from 2-3 independent experiments. Unpaired student’s t-test. **H)** YRNA qPCR of RNAse-treated BALF UF10. Data is expressed as the percentage of remaining YRNA compared to an untreated control. N=6 from 2 independent experiments. Welch’s t-test. I) YRNA qPCR of proteinase K (PK) + RNase-treated BALF UF10. Data is expressed as the percentage of remaining YRNA compared to an RNase-only treated control. N=6 from 2 independent experiments. **J)** YRNA levels from BALF SEC fractions F1-4 in PBS and OVA-challenged mice. Data is expressed as the ΔCt fold change compared to PBS-challenged mice. Unpaired student’s t-test. N=4+ from two independent experiments. An RNY1 value (OVA) was excluded as an outlier using the ROUT outlier test. **K)** YRNA levels from BALF UF10 in PBS and OVA-challenged mice. Data is expressed as the ΔCt fold change compared to PBS-challenged mice. Unpaired student’s t-test. N=4-6 from 2 independent experiments. For all panels: *= P ≤ 0.05, **= P ≤ 0.01, ***= P ≤ 0.001, ****= P ≤ 0.0001. Bars represent the mean +/-standard error of the mean.

Consistent with our results, other studies have shown that YRNAs are present in EVs and protein complexes.^34,45^ However, it is not known how YRNA levels change within these compartments during inflammation. Therefore, we wanted to determine how RNY1 levels in BALF UF100 (EV) and UF10 (RNP) fractions change during airway inflammation. There was a significant ∼5-fold increase in EV RNY1 in OVA-challenged mice **(Fig 4J).** In RNPs, there was also an ∼5-fold increase in RNY1 **(Fig 4K)**. These data show that EV and RNP RNY1 contribute to the total RNY1 level increase during inflammation.

### BALF extracellular vesicle and ribonucleoprotein fractions instruct different macrophage activation programs

Extracellular particles (EPs) from biofluids can provide important instructing signals to recipient cells during immune responses.^48–51^ However, how different EPs from the same complex biofluid alter cellular states is poorly understood. Macrophages are an ideal model system to investigate complex cellular responses to extracellular particles, as they are phenotypically plastic cells that adopt diverse activation states in response to extracellular stimuli from the local microenvironment.^52^ To determine the immunological activity of EVs and RNPs isolated from lung fluid during inflammation, we treated bone marrow derived macrophages (BMDMs) with UF100 and UF10 fractions from BALF after OVA challenge **(Fig 5A)**. We then surveyed a broad panel of macrophage activation genes by qPCR. BMDMs treated with the EV-containing UF100 fraction showed the largest increase in *Arg1* expression, with this gene increasing between 100-1,000 fold after treatment **(Fig 5B)**. There were also statistically significant increases in *Il6, Il10,* and the chitinase-like protein *Ym1* in EV-treated cells. BMDMs treated with the RNP-containing UF10 fraction isolated from the same starting volume of BALF also showed induction of *Arg1 and Il6,* though this treatment induced relatively less *Arg1* and more *Il6* than the EV fraction **(Fig 5C)**. Macrophage treated with the RNP fraction also induced expression of genes that were not induced by the EV fraction including *Tnfa*, *Il1b*, and the interferon responsive gene *Isg15* and failed to induce either *Ym1* or *Il10*. Together, these findings demonstrate that BALF EV and RNP containing fractions are unique signaling compartments with different capacities for instructing macrophage transcription during inflammation.

**Figure 5:**
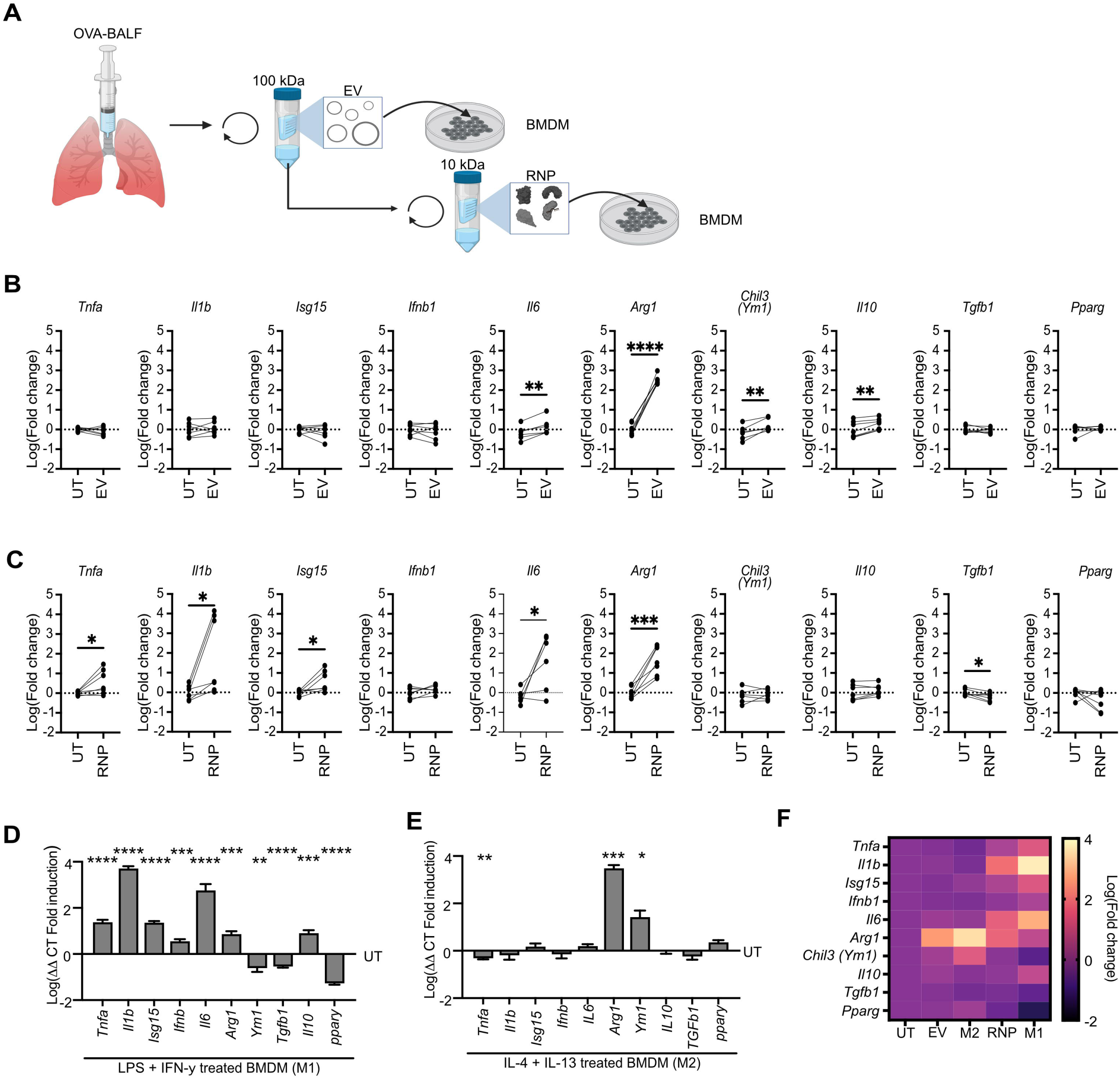
BALF EV RNY1 regulates alternative activation of macrophages during inflammation. **A)** Schema for treating BMDMs with EV and RNP-containing fractions. **B,C)** BMDM gene expression after treatment with EV **(B)** and RNP **(C)** containing fractions determined via qPCR. Each dot represents the fold change in gene expression compared to the untreated control average. Treated values are paired to untreated control BMDMs derived from the same murine bone marrow. Paired student’s t-test. N = 5-7 from three independent experiments. **D,E)** BMDM gene expression after treatment with LPS+IFN-γ(M1) **(D)** and IL-4+IL-13 (M2) **(E)**. Data is expressed as the mean fold change normalized to untreated control BMDMs derived from the same murine bone marrow. Each bar represents the mean gene expression + the standard error of the mean. One sample student’s t-test with a Benjamini/Hochberg correction for multiple comparisons. N = 4-7 from 2-3 independent experiments. **F)** Heatmap depicting BMDM gene expression after treatment with BALF RNP, BALF EV, LPS+ IFN-γ (M1), and IL-4+IL-13 (M2) via qPCR. Each square represents the mean gene expression normalized to an untreated control derived from the same murine bone marrow. N = 4-7 from three independent experiments. For all panels: *= P ≤ 0.05, **= P ≤ 0.01, ***= P ≤ 0.001, ****= P ≤ 0.0001. All Ct values are normalized to the housekeeping gene beta actin.

Given that macrophages treated with the RNP fraction induced expression of more pro-inflammatory genes while macrophages treated with the EV fraction induced expression of genes characteristic of an alternative activation program, we next compared these gene expression signatures with BMDMs activated by a classical pro-inflammatory M1-like stimulus (LPS+IFN-γ) and an alternatively activating M2-like stimulus (IL-4+IL-13) **(Fig 5D,E)**. EV-treated BMDMs showed a strong M2 signature, as both stimulation conditions induced large magnitude increases in *Arg1* and modest increases in *Ym1*. However, BMDMs treated with EVs also induced expression of *Il6* and *Il10* which was not observed in M2 macrophages. RNP-treated BMDMs showed a partial M1 signature, as both RNPs and M1 macrophages upregulated *Tfna, Il1b, Isg15, Il6, and Arg1*, while downregulating *Tgfb1*. However, M1 macrophages also upregulated *Ifnb* and *Il10* while downregulating *Ym1* and *Ppary*, which was not observed in RNP-treated BMDMs **(Fig 5F)**. These results demonstrate that BALF EVs and RNPs induce a gene expression program in macrophages that is related to but distinct from traditional BMDM polarization via isolated pathogen motifs and cytokines.

### RNY1 promotes EV-mediated alternative activation of macrophages by lung fluid

Although RNA levels are dynamic in the extracellular space during many disease processes^2,53^ the function of specific exRNAs is poorly defined. To test how exRNY1 contributes to the function of EVs and RNPs in airway fluid during inflammation, we obtained mice with a CRISPR-mediated knockout of RNY1 **(Fig 6A)**. Mice with both alleles of the RNY1 locus knocked out (*RNY1^-/-^*) were born at normal mendelian frequencies and had no grossly observed phenotypic defects (data not shown). As anticipated, *RNY1^-/-^* mice lacked expression of RNY1 in lung tissue and exRNY1 in BALF **(Fig 6B)**. During OVA-induced airway inflammation, *RNY1^-/-^*mice had normal recruitment of eosinophils, neutrophils, CD4^+^ T cells, and B cells to the lung, indicating that the ability to mount an inflammatory response to allergen challenge remained intact despite systemic loss of RNY1 expression **(Fig 6C).** However, when BMDMs were treated with equal numbers of EVs isolated by ultrafiltration from *RNY1^+/+^* and *RNY1^-/-^* BALF during OVA airway inflammation, macrophages induced significantly less *Arg1* and *Ym1* expression **(Fig 6D)**. No change in *Il6* or *Il10* gene expression was observed. These results demonstrate that EV-mediated induction of M2 genes is partially RNY1 dependent in macrophages. In contrast, when BMDMs were treated with equal concentrations of protein isolated by ultrafiltration from *RNY1^+/+^* and *RNY1^-/-^* inflamed BALF, there was no change in the RNP-inducible genes **(Fig 6E)**. These results demonstrate that RNY1 selectively contributes to airway EV-mediated macrophage activation as part of an extracellular immunologic signaling axis.

**Figure 6:**
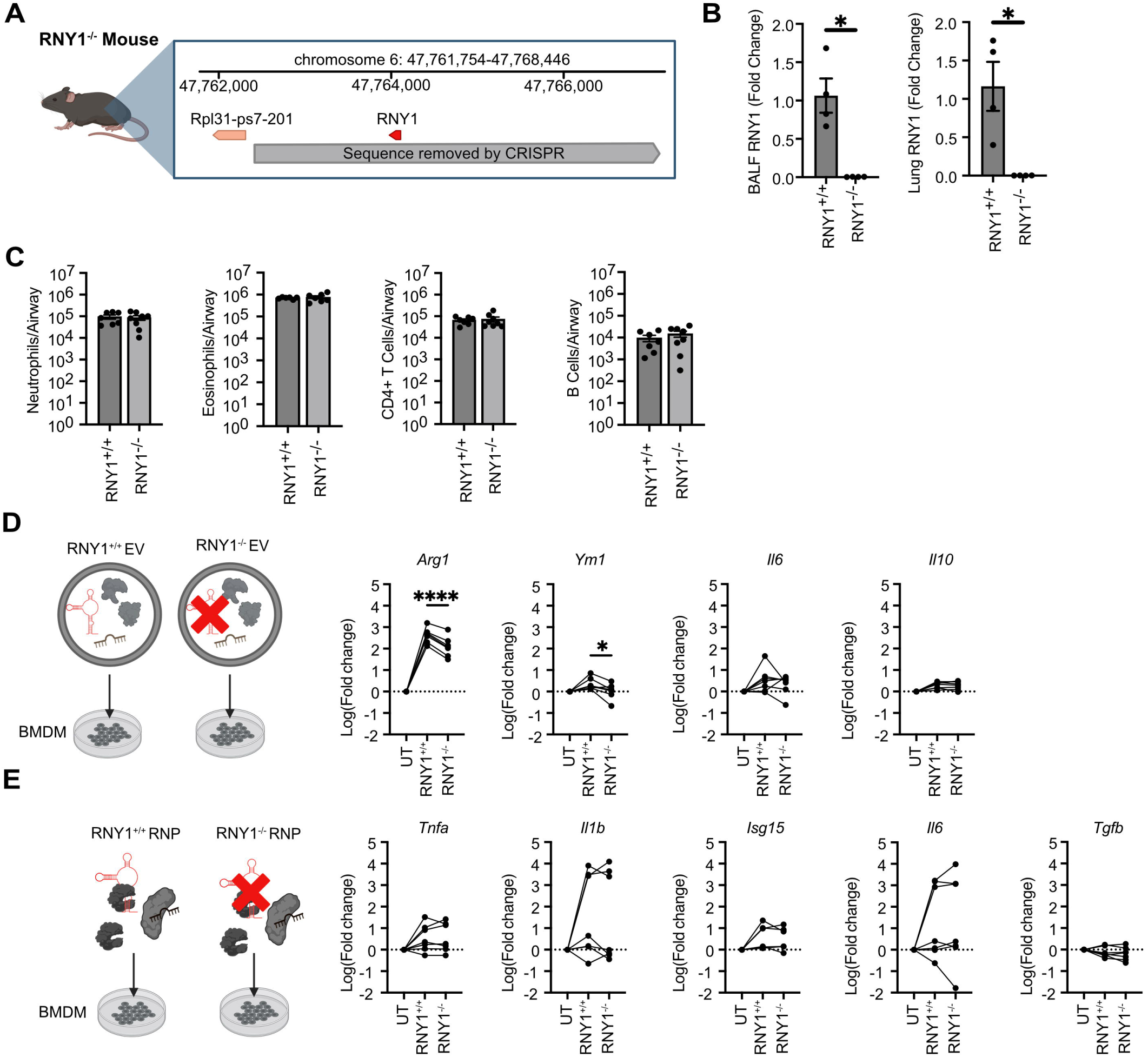
RNY1 regulates BALF EV alternative activation of macrophages during inflammation. **A)** Chromosomal region of CRISPR deletion in *RNY1^-^*^/-^ mice. **B)** RNY1 qPCR of BALF (left) and lung tissue (right) derived from *RNY1^+/+^* and *RNY1^-/-^* mice. Data is expressed as the mean RNY1 fold change compared to the average *RNY1^+/+^* control value. Each bar represents the mean +/-the standard error of the mean. Welch’s t-test. N=4 from 2 independent experiments. **C)** BALF neutrophil, eosinophil, CD4^+^ T cell, and B cell counts from OVA-challenged *RNY1^+/+^* or *RNY1^-/-^*mice determined via flow cytometry. Welch’s t-test. N=6-8 from 2 independent experiments. **D,E)** BMDM gene expression after treatment with EV **(D)** and RNP **(E)** containing fractions derived from *RNY1^+/+^* or *RNY1^-/-^* mice. Data is expressed as the mean log fold change normalized to an untreated control. Treated values are paired to untreated control BMDMs derived from the same murine bone marrow. Paired student’s t-test. N = 5-7 from three independent experiments. For all panels: *= P ≤ 0.05, **= P ≤ 0.01, ***= P ≤ 0.001, ****= P ≤ 0.0001. All Ct values are normalized to the housekeeping gene beta actin.

## Discussion

Although cellular secretion of exRNAs is regulated by inflammatory stimuli,^17^ exRNA dynamics during tissue inflammation is incompletely understood. Our work demonstrates that levels of RNY1 increase ∼8 fold during allergen-induced airway inflammation. The magnitude of RNY1 increase is much larger than that of RNY3, which did not change significantly in bulk biofluid. Importantly, this increase in exRNY1 occurred while cellular levels of RNY1 decreased in the lung, which was not observed for other common Polymerase III transcripts and suggests the possibility of selective YRNA secretion. In addition, our work traces YRNA levels over time and demonstrates that exRNY1 peaks then quickly declines after the inciting inflammatory stimulus. We also observed a strong correlation of neutrophils with exRNY1 levels in the lung. Although eosinophils are the hallmark cells of allergen-induced inflammation, neutrophilic inflammation is common in asthma subsets that are often severe and refractory to current treatments, making identifying novel markers and drivers of neutrophilic lung inflammation in this disease particularly important.^54^

exRNAs can exist outside the cell as free RNAs, as EV cargos, and in protein complexes. Both in cell culture supernatants and blood, prior work has identified exYRNAs in EVs and RNPs.^10,34,35,37,45,55–57^ Our study demonstrates that while RNY3 is only protected in airway fluid by EVs, RNY1 is protected in both EVs and RNPs. These data suggest that the extracellular secretion and/or function of the two members of this small noncoding RNA family may be unique. In addition, although this work adds to prior findings showing that select exYRNA species can be increased during immune responses and diseases,^34,45^ we have found that the relative distribution of YRNAs between EVs and RNPs does not change during airway inflammation. Future work is needed to determine whether the subset of EV or type of protein complex that carries YRNAs changes with inflammation.

Given that exRNAs can be partitioned into multiple extracellular particles, we next wanted to determine how compartmentalization may impact function. Recent literature has shown that EVs and their RNA cargos can provide instructing signals to polarize macrophages to M1-like and M2-like phenotypes.^58,59^ The polarized activation of macrophages by both pathogen-derived and endogenous signals is critical for host defense, tissue damage during inflammation, and reparative functions.^60^ Using a model of allergen-induced inflammation, we found that airway EV fractions upregulated genes more characteristic of M2-like macrophages, which is consistent with the overall type 2 inflammatory response observed in this mouse model of asthma.^61^ In contrast, the airway fluid RNP fractions from this sterile model of inflammation upregulated gene sets with M1-like polarization. These data support the idea that EVs and RNPs instruct macrophages with different, and even contrasting, transcriptional programs. In asthma, the balance of these two immune compartments could be important in mediating the disease endotype and may be significant as patients with less type 2 inflammation have poorly understood pathophysiology. Finally, these observations reflect the complexity of immune regulation within inflamed tissue and the potential of extracellular particles to mediate that inflammation.

To test the role of YRNAs in airway inflammation, we took a genetic approach that allowed us to remove RNY1 from an endogenous biofluid. Interestingly, RNY1 loss attenuated macrophage reprogramming via EVs, but not RNPs. Additionally, RNY1 loss in BALF EVs affected only a subset of genes, as *Arg1* and *Ym1* expression decreased but *Il6* and *Il10* expression did not change in target cells. This work begins to dissect molecular mechanisms of how EVs can function as multimodal intercellular signaling platforms during inflammatory responses within a complex microenvironment, adding to evidence that exRNAs can modulate immune cell responses during type II airway inflammation.^62,63^ These findings align with prior reports that exYRNAs instruct immunological functions in recipient cells such as cytokine secretion, apoptosis, and autoimmune inflammation.^36,37,39,64^ Our study builds on these observations by using an RNY1-free system to show that expression of this YRNA is necessary for EVs from lung fluid to induce robust M2 macrophage gene expression. This adds to prior studies showing the programming of macrophage responses by EV YRNA, and points toward unique features of this programming depending on YRNA species, cell of origin, and pathologic context. Together this work provides a foundation for future studies to investigate the signaling mechanism by which exYRNAs program macrophages in inflammation.

Finally, there are several important limitations to our work. This study uses crude preparations of lung biofluid due to practical limitations in the amount of material we can recover. With continued improvements in technologies to purify EVs and other extracellular particles, there will be opportunities to pinpoint which RNY1-containing EV subpopulations are immunologically active in airway fluid. In addition, our experimental strategies do not identify which form of RNY1 is important in inflammation. Our qPCRs capture both full-length and fragmented YRNAs, and our RNY1 knockout removes both forms from the mouse. Therefore, other approaches will be needed to determine if there are differences in the dynamics and function of full length and fragmented YRNAs. exRNAs have expanding roles as mediators of intercellular communication, including during immune responses.^17^ This work identifies exRNY1 in both EV and RNP compartments within airway lining fluid and establishes this YRNA as a modulator of immune cell gene expression.

## Abbreviations

EV: Extracellular vesicle
EP: extracellular particle
RNP: Ribonucleoprotein particle
OVA: ovalbumin
HDM: House dust mite
BALF: Bronchoalveolar lavage fluid
UF100: ultrafiltrated extracellular material >100 kDa
UF10: ultrafiltrated extracellular material <100 kDa and >10 kDa
Tx: Triton X-100
PK: Proteinase K
SEC: size exclusion chromatography

## Acknowledgements

We thank Vanderbilt’s Center for Extracellular Vesicle Research (CEVR), Cell and Developmental Biology Core (CBD) for use of their equipment. We thank the Vanderbilt University Medical Center Division of Animal Care for their mouse procurement, husbandry, and healthcare. We thank Dr. Wenhan Zhu for use of the 384-well qPCR machine. Figure graphics were designed using Biorender (www.Biorender.com)

## Data availability statement

The data supporting the findings of this study are available from the corresponding author upon request.

## Author Contributions

**Cherie E Saffold:** Conceptualization (lead), methodology (lead), data acquisition (lead), data analysis (lead), data visualization (lead), writing – original draft (lead), writing – review and editing (lead). **Antiana Richardson:** methodology (support), data acquisition (support), writing – review and editing (support). **Heather H Pua:** Conceptualization (lead), methodology (lead) funding acquisition (lead), resources (lead), project administration (lead), writing – original draft (lead), writing – review and editing (lead)

