## Supplemental Figures for "RNY1 partitions into extracellular vesicles and ribonucleoprotein particles during airway inflammation to regulate macrophage programming"

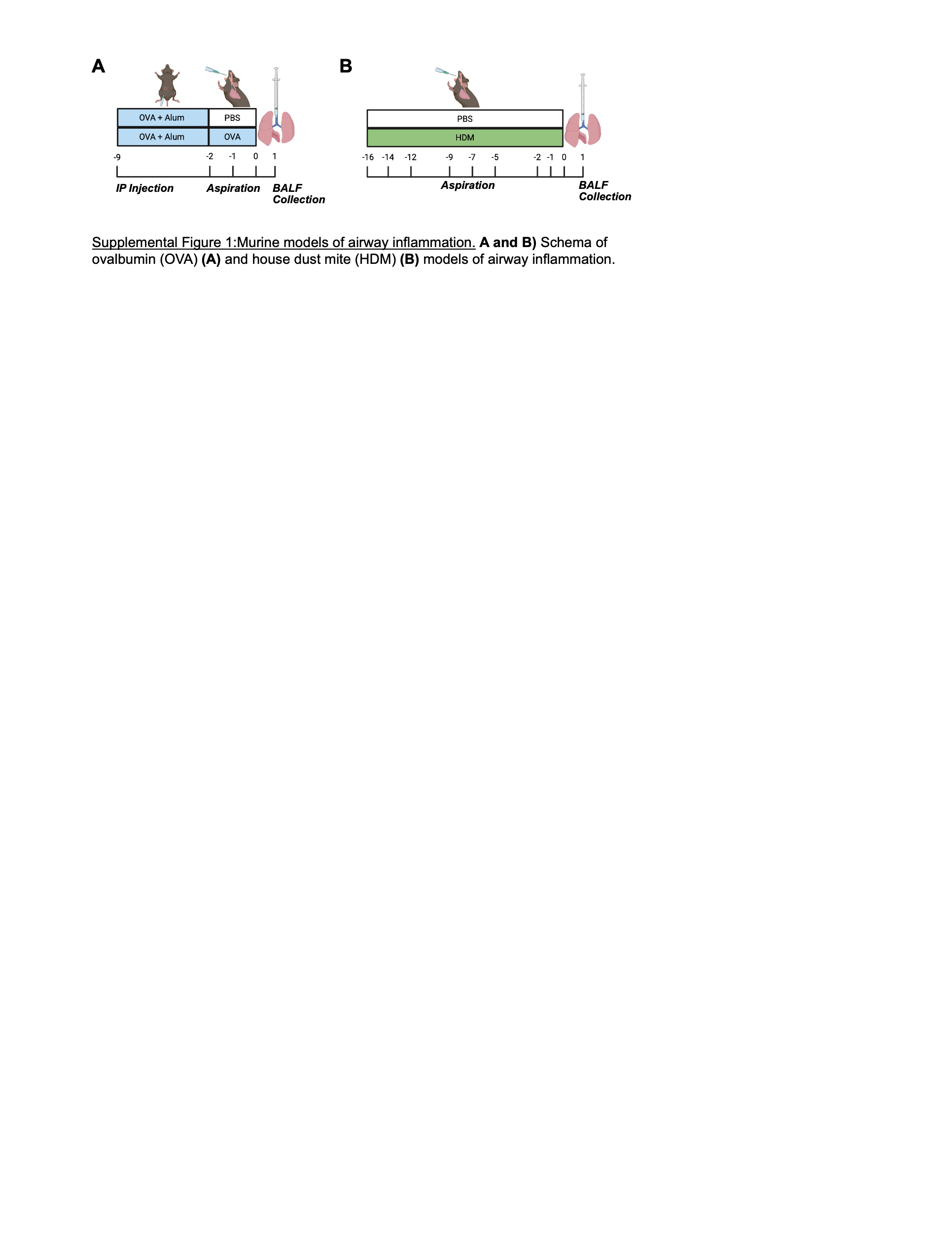

**Supplemental Figure 1:** Murine models of airway inflammation. A and B) Schema of ovalbumin (OVA) (A) and house dust mite (HDM) (B) models of airway inflammation.

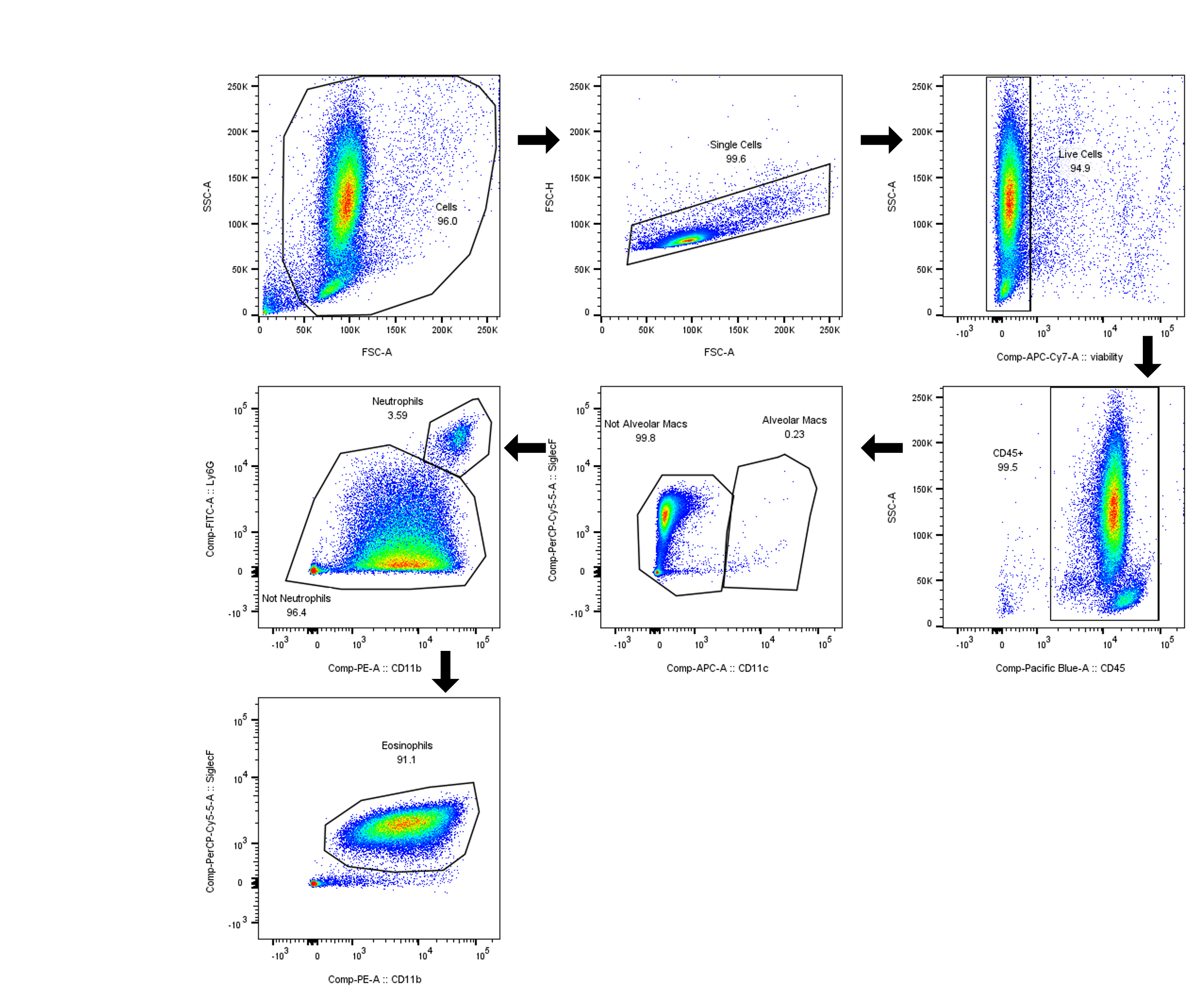

**Supplemental Figure 2:** Flow cytometry gating strategy for alveolar macrophages, neutrophils, and eosinophils in BALF.

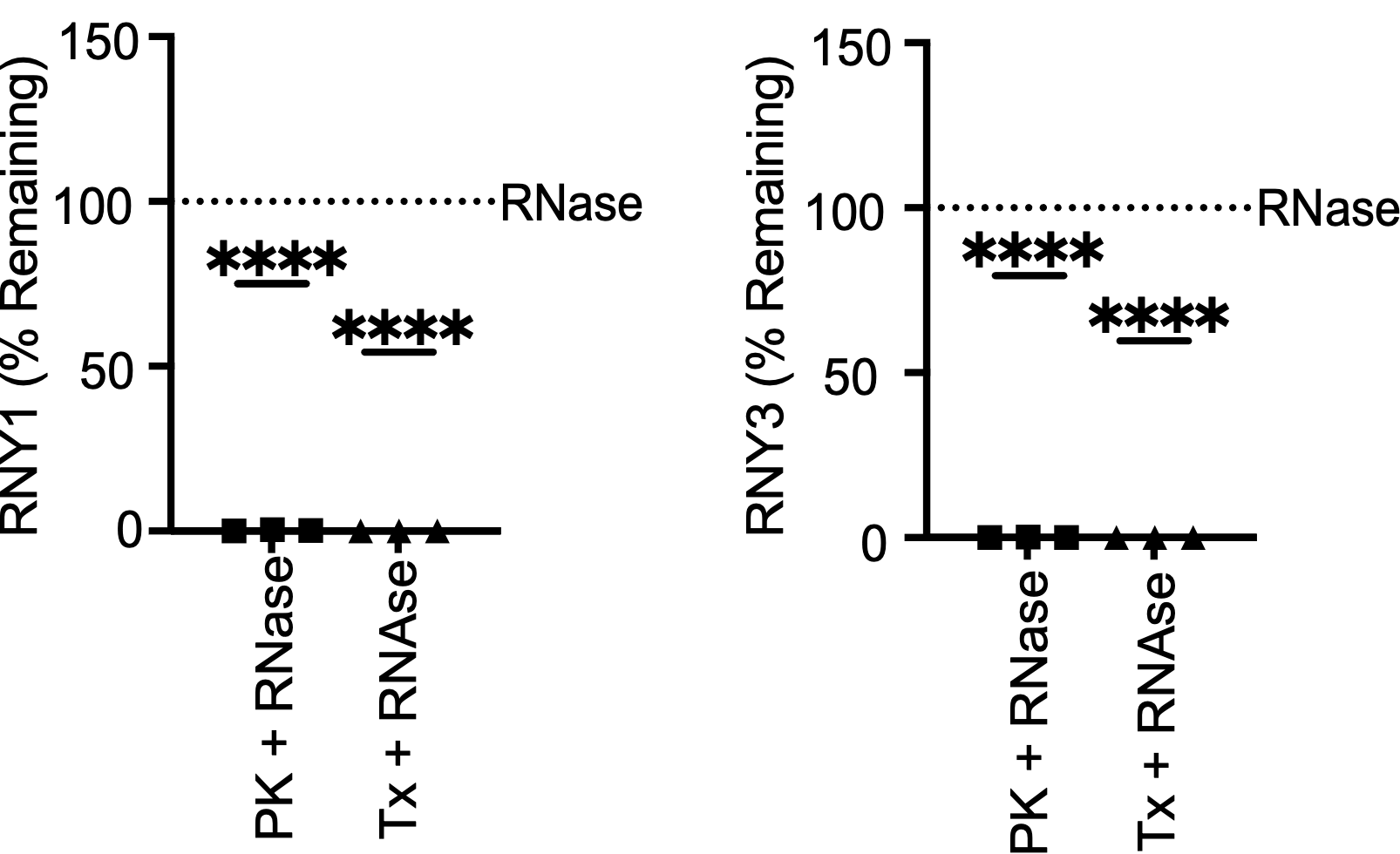

**Supplemental Figure 3:** RNAse sensitivity assays on extracted BALF RNA. YRNA qPCR of RNAse + PK and RNase + Tx treated BALF RNA pre-extracted with phenol-chloroform. Data is expressed as the percentage of remaining YRNA compared to an untreated control. One sample student's t-test. N=3 from 2 independent experiments.

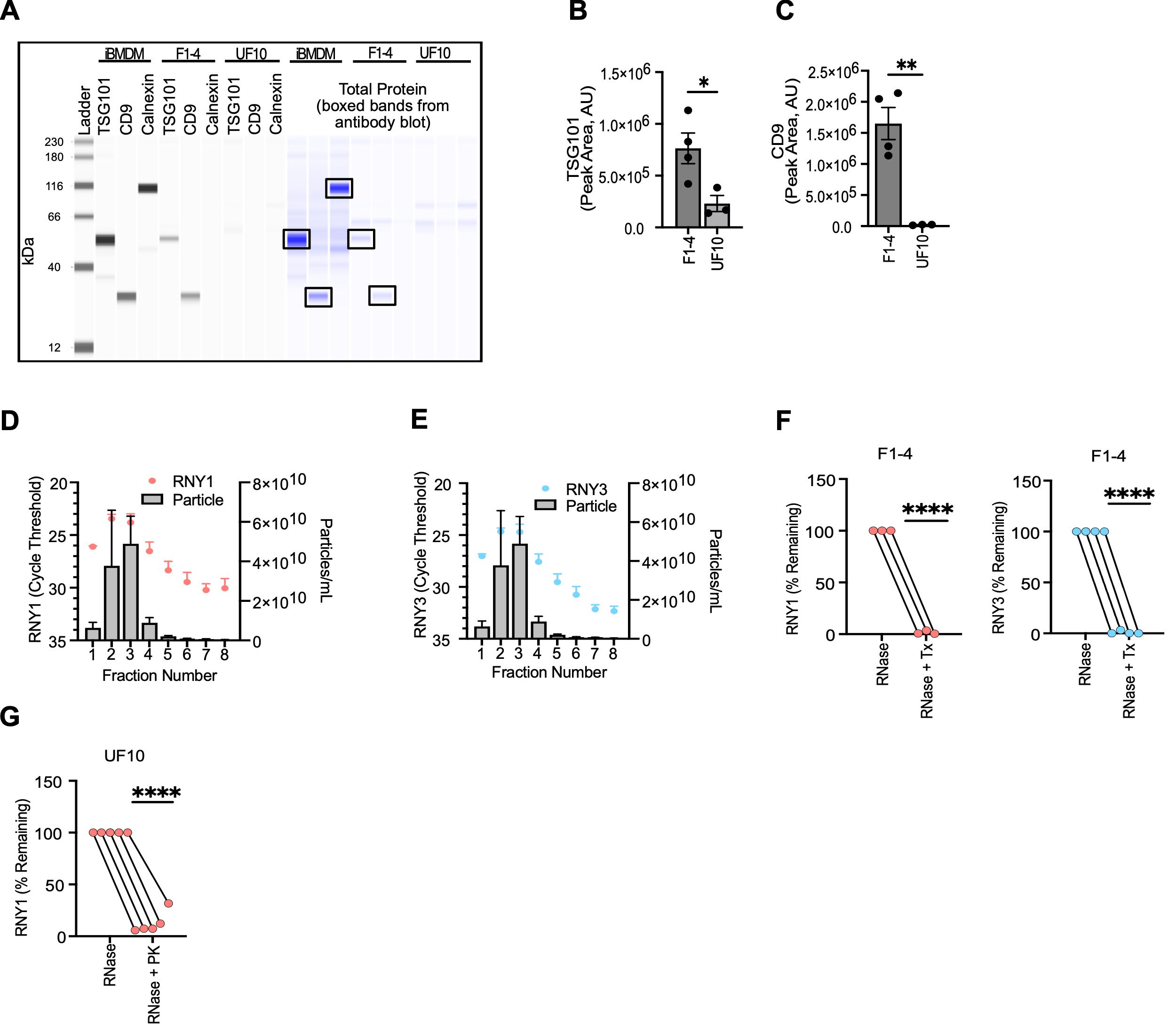

**Supplemental Figure 4:** YRNA partitioning in extracellular vesicles and ribonucleoproteins in PBS-challenged control mice. A) Western blot of SEC purified EV fractions 1-4 and RNPs captured in UF10. Western blot was performed using a capillary-based electrophoresis system virtual, blot-like representations of chemiluminescent signals for each antibody, or a biotin label for total protein. Immortalized bone marrow derived macrophages were used as a positive control for calnexin detection. Western blot is representative of 3-4 samples from 2 independent experiments. B,C) Quantification of chemiluminescent detection of TSG101 (B) and CD9 (C). Area peaks were normalized to total protein detection in each lane. N=3-4 from 2 independent experiments. Unpaired student's t-test. D,E) RNY1 (D) and RNY3 (E) qPCR Ct values and particle counts from SEC fractionated BALF UF100 from PBS-challenged mice. N=3 from 2 independent experiments. F) YRNA qPCR of Triton X-100 (Tx) + RNAse treated BALF SEC fractions 1-4 from PBS-challenged mice. Data is expressed as the percentage of remaining YRNA compared to an RNAse only treated control. One sample student's t-test. N = 4 from 2 independent experiments. One RNY1 value was removed as an outlier using the ROUT outlier test. G) RNY1 qPCR of proteinase K (PK) + RNAse treated BALF UF10 from PBS challenged mice. Data is expressed as the percentage of remaining YRNA compared to an RNAse only treated control. N=5 from 2 independent experiments. One sample student's t-test. For all panels: *= P ≤ 0.05, **= P ≤ 0.01, ***= P ≤ 0.001, ****= P ≤ 0.0001. Bars represent the mean +/- standard error of the mean.
